# Did I imagine that? The functional role of paracingulate cortex in reality monitoring

**DOI:** 10.1101/2020.05.19.103572

**Authors:** JR Garrison, F Saviola, E Morgenroth, H Barker, Michael Lührs, JS Simons, C Fernyhough, P Allen

## Abstract

Reality monitoring describes our ability to distinguish between internally and externally generated experiences. Individuals show significant variation in this ability and impaired reality monitoring has been linked to the experience of hallucinations. We undertook two studies to investigate the association between reality monitoring and morphology of the paracingulate region of medial prefrontal cortex. In Study 1 we compared reality monitoring accuracy and functional connectivity within paracingulate cortex in groups of healthy controls (N=20) and patients with schizophrenia and hallucinations (N=19). Controls showed greater reality monitoring accuracy that was associated with resting-state functional connectivity between paracingulate, precuneus and occipital cortices, while reality monitoring in patients was associated with more lateral functional connectivity. In Study 2 we used real-time fMRI neurofeedback to obtain causal evidence for the role of the paracingulate cortex in reality monitoring. Healthy individuals received Active feedback from paracingulate cortex (N=21) or Sham feedback based on randomised signal (N=18). Active-group participants showed a specific behavioural effect of improved reality monitoring for Imagined items, as well as increases in both activity within the paracingulate region, and its posterior functional connectivity with precuneus and lateral parietal cortices, and occipital cortex.

Our findings suggest reality monitoring in healthy individuals is causally supported by a paracingulate mediated flexible network including the precuneus. Network connectivity can be enhanced using neurofeedback and tracks with improved reality monitoring ability. In contrast, patients with schizophrenia may utilise a distinct and more lateral network which may explain observed sub-optimal reality monitoring accuracy, contributing to the experience of hallucinations.

**Significance Statement:** Reality monitoring refers to our ability to distinguish imagination from our experiences in the outside world, and is linked both to hallucinations in schizophrenia as well as to the morphology of paracingulate cortex area of the brain. Here, we revealed less paracingulate involvement in the functional reality monitoring networks in patients with schizophrenia compared to healthy individuals. Thereafter, we used real-time fMRI neurofeedback to show that healthy individuals can learn to upregulate brain activity within the paracingulate cortex, with this resulting in both improved reality monitoring ability and changes in paracingulate functional connectivity. This suggests that paracingulate cortex activity and connectivity play a causal role in reality monitoring, with implications for both the understanding and treatment of hallucinations.

## Introduction

We have all had the experience of being unable to remember whether an event in our past really happened, or was something we imagined. Reality monitoring refers to the cognitive processes we use to distinguish internally generated experiences from those perceived in the external world (1). Healthy individuals show a high level of variation in this ability (2), and impairments in reality monitoring have been linked to the experience of hallucinations in patients with schizophrenia (3). Reality monitoring ability, as measured by recollection of the perceived vs. imagined aspects of context, is associated with functional activity and connectivity within the medial prefrontal cortex (mPFC) (4–6), consistent with the role of cortical midline structures more generally in self-referential processing (7–9). This is further supported by evidence linking the structural morphology of the paracingulate sulcus (PCS) region of the mPFC to both reality-monitoring ability in healthy individuals (2), and to the experience of hallucinations in patients with schizophrenia (10, 11). While there is currently no causal evidence linking reality monitoring ability to functionality either generally within the mPFC, or more specifically within paracingulate cortex, it is suggested that altered activity and functional connectivity within this brain region may underlie the behavioural reality monitoring impairments observed in schizophrenia patients, contributing to their experience of hallucinations (12–14).

We conducted two Studies to seek a functional explanation for the link between reality monitoring ability, activity and connectivity within the paracingulate region of the mPFC. In the first of these, we investigated the differential functional networks associated with reality monitoring in patients with schizophrenia and demographically matched healthy controls looking for differences within the paracingulate cortex and other cortical midline regions of the brain which have been associated with self-referential processing (7, 15). We built on these findings in our second study using real-time fMRI neurofeedback (henceforth ‘fMRI neurofeedback’) to gain causal evidence for the role of paracingulate cortex in reality monitoring in healthy individuals, predicting that successful up-regulation of regional mPFC activity would lead to improvements in reality monitoring accuracy and changes in functional connectivity within a similar reality monitoring brain network to that identified in healthy subjects in Study 1.

In both studies, we focused specifically on recollection of the source of imagined information, as hallucinations in schizophrenia are linked both to the failure to recognise the source of self-generated/internal content (16, 17) as well as to enhanced externalising bias, by which imagined information is more likely to be recognised as having been perceived, than perceived information is to have been imagined (reviewed in (13)). We also focused on functional connectivity differences associated with two previously defined networks which both encompass regions of paracingulate cortex – the Default Mode and Somatosensory Networks (DMN & SSN) (18). The choice of these networks was key to our first study. The DMN includes anterior paracingulate cortex /precuneus, lateral parietal cortices and the hippocampus (18), and is active when the brain is at rest (19). Changes in activity within the DMN, and especially in its core medial areas, are associated with a wide variety of tasks which involve self-referential processing (7, 15). For example, activity within the DMN during task-related reality monitoring judgements was found to be more positive (i.e. reduced task-specific deactivation) in healthy individuals when compared with an external source monitoring task (4). In contrast, the SSN includes posterior paracingulate cortex, motor and somatosensory cortices, and lateral areas around the superior temporal gyrus (STG) including auditory cortex (18). The STG is also active during self-referential processing tasks (20, 21), with auditory regions of the STG active during state studies of auditory verbal hallucinations (22). Changes in mPFC to left STG functional connectivity have been linked to variable self-other reality monitoring judgements in schizophrenia (23), while hallucinations in schizophrenia have been associated with dysfunctional functional connectivity of left STG, as well as more dysconnectivity within the DMN and its interaction with other resting state networks (24).

In Study 1, we used an established reality monitoring task paradigm to compare reality monitoring accuracy for imagined items and functional connectivity between healthy controls and patients with schizophrenia that experience hallucinations. Evidence suggests that reality monitoring accuracy is associated with activity and functional connectivity within the mPFC (4, 23) as well as more posterior medial brain regions implicated in self-referential processing including the precuneus (5, 7, 8, 25). We therefore used exploratory Independent Component Analysis (ICA) to restrict our investigation into whether group differences in functional connectivity in midline cortical brain regions within the DMN and SSN related to impairments in patients’ behavioural reality monitoring performance. We predicted that, relative to healthy controls, patients with schizophrenia would show impaired reality monitoring performance for Imagined items that is associated with differential functional connectivity in paracingulate brain regions.

Thereafter, in Study 2, and based on our findings from Study 1, we used fMRI neurofeedback to establish causal evidence for the involvement of paracingulate cortex in recollection of the source of imagined information in healthy individuals. Our choice of fMRI neurofeedback as a technique follows its successful recent deployment in a range of studies where individuals have been trained to self-regulate neural activity in brain regions and networks thought to underlie certain behaviours (26), cognitive functions (27, 28) and psychiatric symptoms (29, 30). As fMRI neurofeedback allows individuals to monitor and self-regulate their own brain activity in real time (31), training individuals to regulate neural activity in particular brain regions or networks enables causal inferences to be made about that region’s involvement in a certain behaviour or function. Moreover, if a region is known to be involved in potentially distressing atypical experiences, such as hallucinations, then altering activity in that region may have therapeutic benefits.

Using a sham-controlled between-subjects design, we investigated whether healthy individuals could learn to up-regulate paracingulate cortex activity via fMRI neurofeedback training, and whether successful up-regulation would lead to improvement in reality monitoring accuracy for imagined information, i.e. comparing performance on a reality monitoring task at pre-and post fMRI neurofeedback training time-points. We also explored whether veridical fMRI neurofeedback would result in functional connectivity changes between the paracingulate cortex and other brain regions. Specifically, we predicted that up-regulation of paracingulate cortical activity would improve reality monitoring accuracy for Imagined items but not general item recognition memory, and that this would be associated with increased functional connectivity within the medial self-referential network identified in healthy individuals in Study 1 and previously associated with self-referential processing.

## Study 1. A comparison of reality monitoring ability and associated functional connectivity between patients with schizophrenia and healthy controls

### Results

#### Reality monitoring performance

Patients with schizophrenia showed impaired Perceived/Imagined reality monitoring performance relative to healthy controls: F(37,1) = 10.823, p = .002, η_p_^2^ = .181. Post hoc testing revealed significantly lower accuracy in patients for recollection of the source of Imagined items (t(37) = 3.042, p = .005, d = 0.980 word-pairs) but not for Seen items (t(37) = 1.411, p = .167, d = 0.451; Figure 1A). However, the group by condition interaction did not reach significance, F(37,1) = 0.990, p = .326, η_p_^2^ = .026.

**Figure 1.**
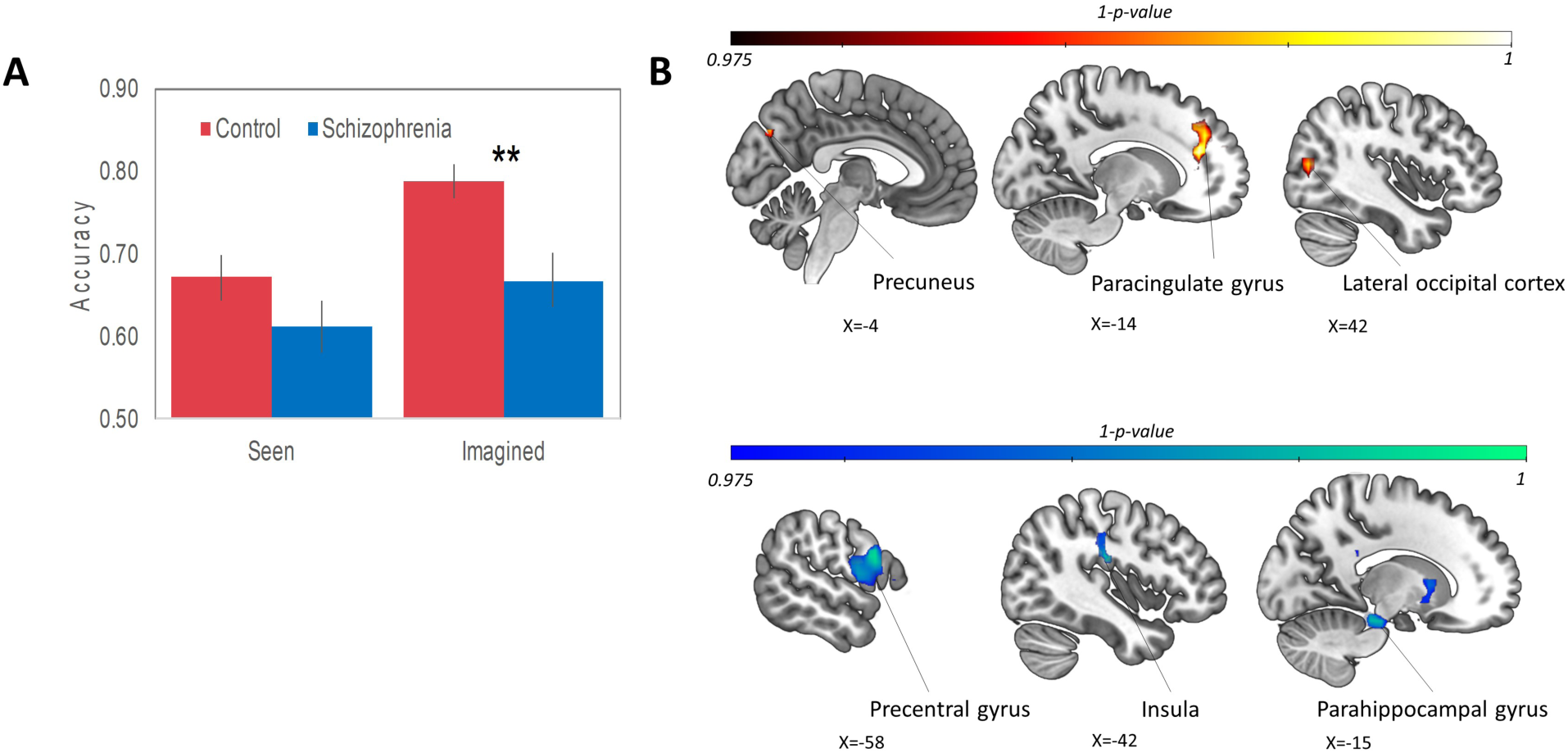
Study 1 results. **A)** Mean accuracy (proportion of accurate responses) by group for Imagined and Seen items where 0.5 represents chance level of accuracy. Nb. All error bars = Standard Error. Patients with schizophrenia (blue, N = 19) show significantly reduced reality monitoring accuracy compared with healthy controls (Red, N = 20) for recollection of the source of Imagined items, but not for the source of previously Seen items (** indicates p < .01). **B)** Brain maps showing functional connectivity interaction effects (group x reality monitoring accuracy) within the DMN in regions where reality monitoring accuracy was positively (upper row) or negatively (lower row) correlated with functional connectivity in healthy controls, thresholded at p < .075 uncorrected for visualisation purposes. Note: Details of these effects and their scatter plots are given in Supplementary Materials Table S2 and Figure S3.

#### Resting state functional connectivity and reality monitoring

Relative to controls, patients with schizophrenia showed both increased and decreased functional connectivity in the SSN component, but there were no significant group differences within the DMN component (see Supplementary Materials Figure S2 and Table S1). We did however observe a significant interaction between group and reality monitoring accuracy for Imagined items on functional connectivity within the DMN Independent Component only (Figure 1B, & Supplementary Materials Table S2 & Figure S3). In the control group there was a significant positive association between reality monitoring accuracy for Imagined items and functional connectivity in the paracingulate gyrus [−14, 38, 20; all coordinates are MNI unless otherwise stated] and precuneus [−6, −74, 36], as well as in the lateral occipital cortex [42, −74, 12], i.e. increased reality monitoring accuracy was associated with increased functional connectivity in these regions. In patients with schizophrenia, however, the association between accuracy for Imagined items and functional connectivity in these regions was negative (Figures 1 and S3). In contrast, in patients with schizophrenia, reality monitoring accuracy was positively correlated with increased functional connectivity in the parahippocampal gyrus [−10, −34, −16], fusiform cortex [42, −38, −32], insula [−42, −22, 24] and precentral gyrus [−58, 10, 24], regions where reality monitoring accuracy was associated with decreased functional connectivity in healthy controls (Figures 1 & S3).

## Study 2. The effect of fMRI neurofeedback targeted on paracingulate cortex activity, on reality monitoring in healthy participants

### Results

#### fMRI Neurofeedback

Analysis of the fMRI data over three neurofeedback training runs revealed a significant group (Active vs. Sham) x Run (Run1, Run2, Run3) interaction in the activity-based paracingulate region of interest (ROI; Blue sphere in Figure 2).

**Figure 2.**
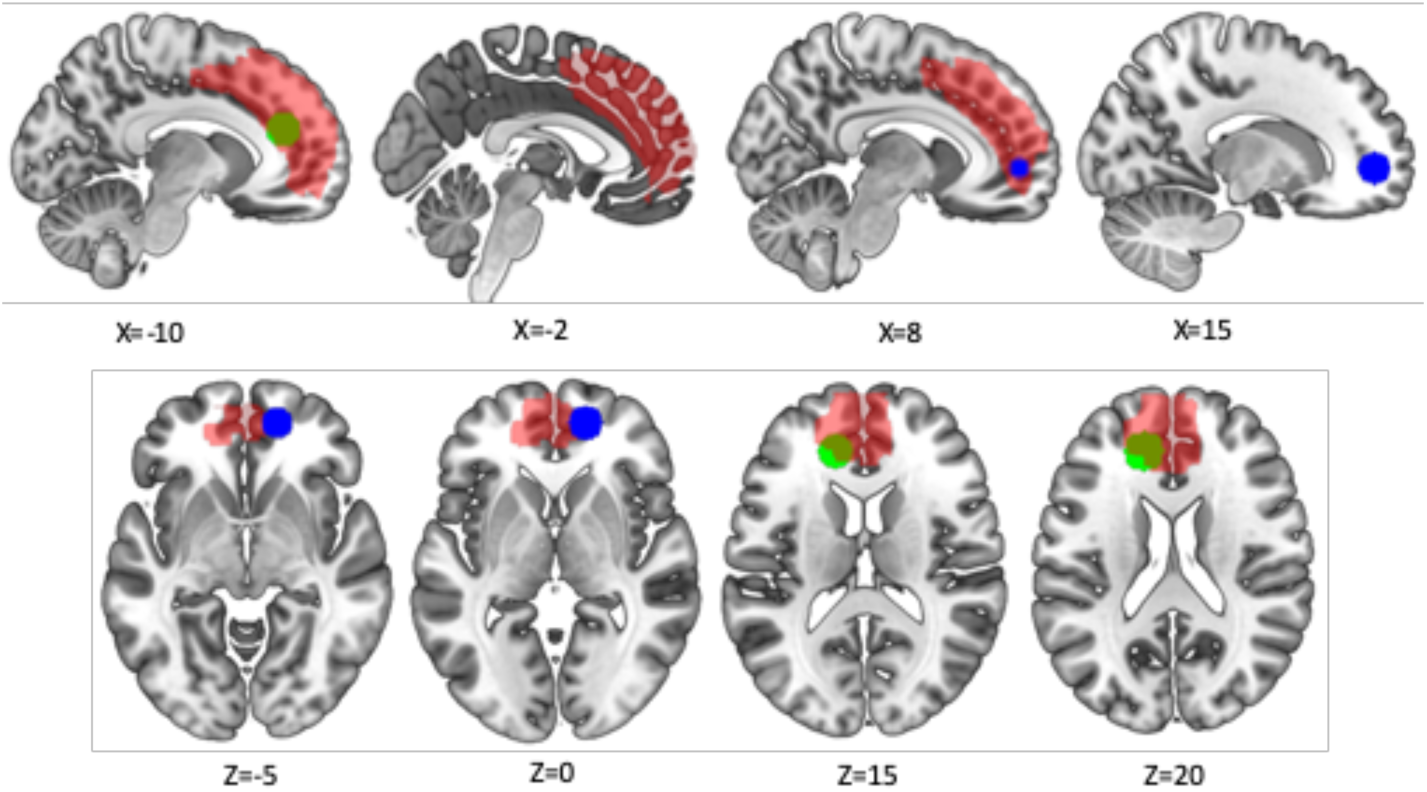
Study 2 fMRI neurofeedback analysis: location of the activity-based paracingulate ROI dysfunctional in schizophrenia and associated with reality monitoring performance (25, 32) [Blue sphere, Talairach: 14, 47, −1], and connectivity ROI based on paracingulate peak of reality monitoring x functional connectivity interaction from Study 1 [Green sphere, −14, 38, 20]. Both ROI’s overlapped with the paracingulate VOI mask across all 39 participants (Red).

Peak activity was observed at [8, 48, −4; Z = 2.79, p_family-wise error (FWE) peak_ = .045, small volume correction (SVC)], which lay within the mean paracingulate voxel of interest (VOI) mask of all 39 participants (Figure 3A).

**Figure 3.**
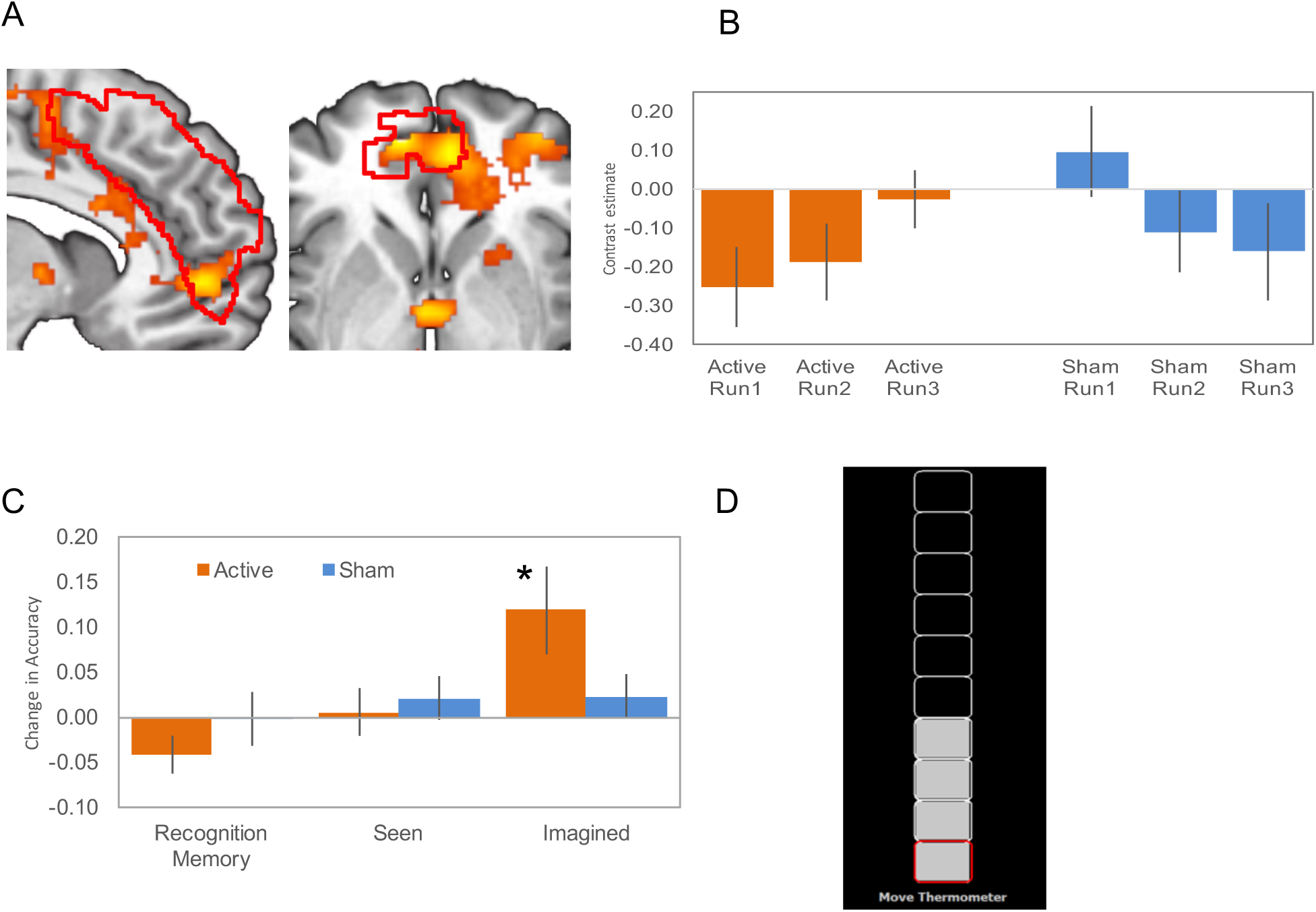
Study 2 fMRI and behavioural results and example neurofeedback gauge (A) fMRI activity showing the group (Active > Sham neurofeedback) by training run (Run1, Run2, Run3) interaction effect, lying within the paracingulate sulcus VOI mask across all 39 participants (outlined in red), images centred on the peak mPFC voxel (8, 48, −4), activity thresholded at p < .075 uncorrected for visualisation purposes. (B) Plot from the peak voxel [8, 48, −4] showing increased activity for the Feedback > Rest contrast across the three neurofeedback runs in the Active group (Orange, N = 21), and decreased activity in the Sham group (Blue, N = 19). (C) Change in reality monitoring and recognition memory behavioural accuracy (Post > Pre-neurofeedback training) in Active and Sham groups (D) Example of the visual neurofeedback gauge interface.

Post-hoc tests showed significantly increased activity in the Active groups across training runs (p_FWE peak_ = .031, Z = 2.82, SVC), whereas the Sham group exhibited a non-significant decrease (p_FWE peak_ = .102, Z = 2.40, SVC; Figure 3B). Thus, participants receiving Active neurofeedback showed differentially increased functional activity within the Paracingulate VOI used for the fMRI neurofeedback training.

#### Reality monitoring

Following neurofeedback training we observed a significant improvement in accurate recollection of the source of Imagined items (Figure 3C). To investigate this, we carried out a mixed (group by session) ANOVA for imagined item accuracy which revealed a main effect of session, F(37,1) = 6.054, p = .019, η_p_^2^ = .141, but not of group, F(37,1) = .074, p = .787, η_p_^2^ = .002. Post-hoc testing revealed a significant improvement in accuracy for Active, t(20) = 2.422, p = .025, d = .528, but not Sham feedback, t(17) = .946, p= .357, d = .223 (Figure 3C). However, the group x session interaction for Imagined items did not reach significance, F(37,1) = 2.700, p =.109, η_p_^2^ = .068. There were no significant pre/post-training neurofeedback effects on accuracy for item recognition, nor for accuracy in the recognition of the source of previously Seen items. There were no significant correlations between the signal change in the peak paracingulate voxel [8, 48, −4] between training Run1 and Run3, and the change in reality monitoring accuracy for recollection of the source of Imagined items in either group (r < .093, p > .485).

#### Resting state functional connectivity

We compared the effect of Active fMRI neurofeedback (post vs. pre) on paracingulate functional connectivity with seed-based analysis based on the reality monitoring interaction peak from Study 1 [−14, 38, 20]. This revealed increased functional connectivity with a single large cluster (p = .009) with peaks in lateral occipital cortex, precuneus and supramarginal gyrus overlapping with SSN activation (Figure 4 & Supplementary Materials Table S3).

**Figure 4.**
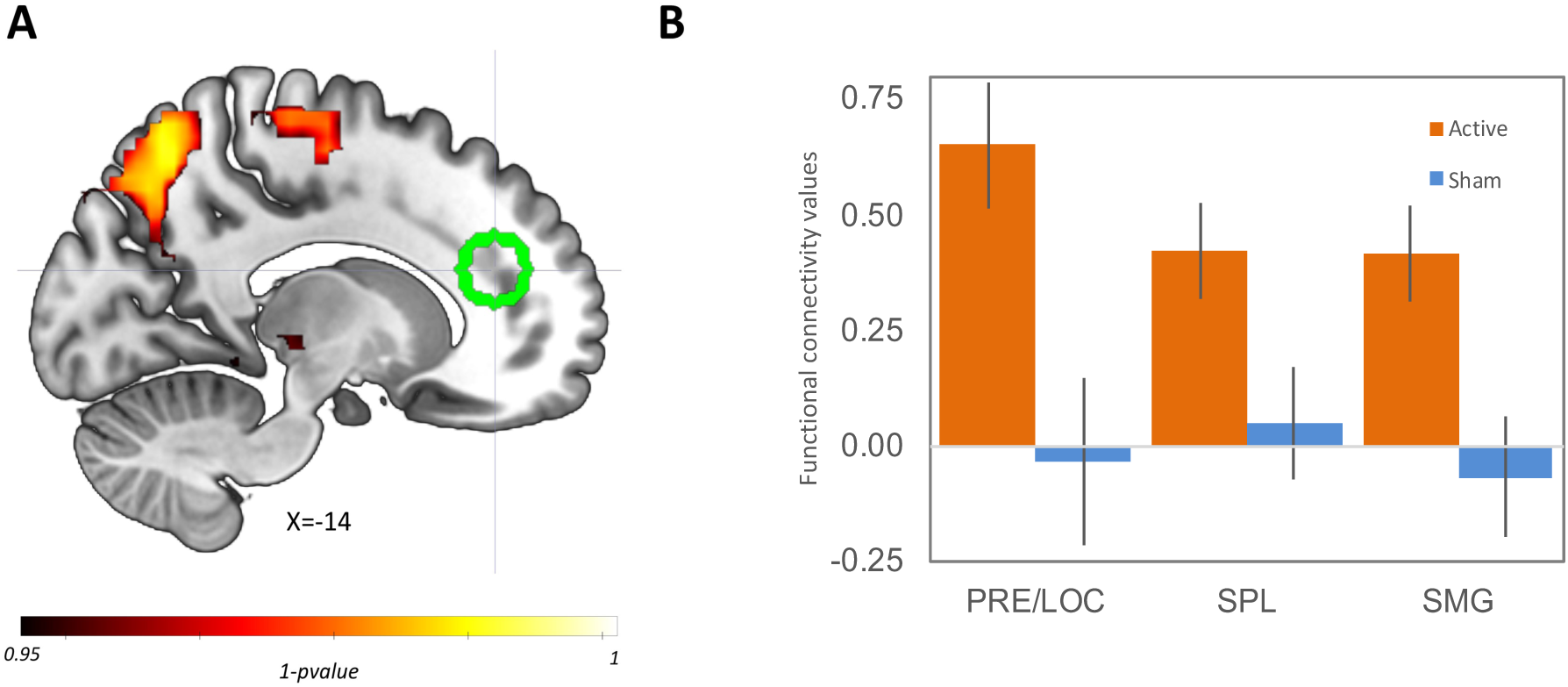
Study 2 functional connectivity results (A) Changes in functional connectivity following Active fMRI neurofeedback associated with the paracingulate seed region (Green sphere, centred on - 14, 38, 20); (B) Group changes in functional connectivity within the three strongest peaks of the Active cluster shown in A, visually indicating the specificity of the effect of fMRI neurofeedback on functional connectivity within these regions, to the Active group alone (see Supplementary Materials Table S3). Note: PRE = Precuneus, LOC = lateral occipital cortex, SPL = superior parietal lobule, SMG = supramarginal gyrus.

A group (Active vs. Sham) by timepoint (pre vs. post fMRI neurofeedback training) interaction also revealed a trend effect for change in functional connectivity with the paracingulate region, showing increased precuneus connectivity in the Active group compared to Sham (p = .07). We found no regions showing significant correlations between changes in functional connectivity and the change in reality monitoring accuracy for Imagined items in the Active group (p < .19).

## Discussion

Our objective for this study was to investigate the functional role of paracingulate cortex in reality monitoring and explore differences in paracingulate cortex connectivity that might contribute to the reality monitoring impairment widely observed in patients with schizophrenia. A particular aim was to seek causal evidence for the role of paracingulate cortex in reality monitoring and explore this region’s wider network functional connectivity. In Study 1, we report differential and more lateral network connectivity underlying reality monitoring in patients with schizophrenia, which may contribute to their behavioural impairment and the experience of hallucinations. Thereafter in Study 2, we demonstrate that reality monitoring accuracy for imagined items in healthy individuals can be improved by fMRI neurofeedback training, suggesting causal involvement of paracingulate cortex. We also show that fMRI neurofeedback training can result in changes in functional connectivity consistent with the reality monitoring network indicated in our first Study.

In line with our hypotheses in Study 1, we found that patients with schizophrenia with hallucinations had lower reality monitoring accuracy for imagined information compared to matched healthy controls, and differential functional connectivity within the SSN Independent Component. Variable reality monitoring accuracy in healthy individuals was specifically associated with differential resting-state functional connectivity in predominantly midline structures of the DMN including both paracingulate and precuneus cortex. Notably, these included the midline cortical regions of paracingulate/cingulate gyri, and the precuneus where activity has previously been associated with self-referential processing (7, 15) as well as lateral occipital cortex. In these three regions, there was a positive association between functional connectivity and reality monitoring in healthy controls and a negative association in patients. This contrasted with reality monitoring accuracy in patients where the associated functional network involved more lateral brain regions including insula and precentral/ inferior frontal gyri. Specifically, in this lateral network including the parahippocampal gyrus, temporal occipital cortex, insula and precentral gyrus, we observed the reverse pattern – that of a positive association between reality monitoring accuracy and functional connectivity in patients and negative in healthy controls. These latter regions have no notable earlier association with self-referential processing perhaps explaining the sub-optimal reality monitoring ability in the patient group. Thus, differences in behavioural reality monitoring accuracy for imagined items between patients with schizophrenia and healthy controls may be explained, at least in part, by modulation of a medially-focused self-referential functional network subserving this ability. Moreover, this finding of impaired reality monitoring in patients with schizophrenia is consistent with earlier behavioural reports (summarised in (13)), and with our work in this same sample of participants showing the reality monitoring impairment to be associated with task specific dysfunctional mPFC activity (12). Our current research thus complements these earlier findings to reveal differences in functional connectivity which may explain the reality monitoring impairment observed in schizophrenia.

Our results from Study 1 thus provide evidence for the involvement of paracingulate cortex in a functional reality monitoring network in healthy individuals, and suggest that the link between PCS morphology, reality monitoring (2) and hallucinations in schizophrenia (10) may be explained by variation in paracingulate functional connectivity. We extended this research using real-time fMRI neurofeedback in our second Study to investigate a causal link between reality monitoring, paracingulate brain activity and wider functional connectivity. We found that participants who received Active neurofeedback from the paracingulate cortex were able to successfully up-regulate activity within this brain region over the course of three neurofeedback training runs undertaken in a single scanning session. Active group participants also demonstrated improved reality monitoring accuracy for imagined items following scanning, and increased functional connectivity between paracingulate cortex (used as a seed in the functional connectivity analysis) and precuneus, occipital and lateral parietal cortices (supramarginal and post central gyri and superior parietal lobule). This indicates there was significant overlap between the brain regions showing enhanced functional connectivity following Active fMRI neurofeedback and those showing a positive group connectivity interaction effect in healthy individuals with reality monitoring accuracy from Study 1 (cingulate/paracingulate, precuneus and lateral occipital cortex). Thus, up-regulation of paracingulate activity following Active fMRI neurofeedback may have resulted in increased functional connectivity within the medial self-referential reality monitoring network in healthy individuals, resulting in improved reality monitoring accuracy for imagined items.

This effect of neurofeedback training appeared specific to recipients in the Active group. Increased paracingulate activity was not observed for participants receiving the Sham neurofeedback signal, and a functional connectivity interaction analysis between the groups (Post vs Pre scanning) showed there to be a trend effect for change in the Active group compared to the Sham within paracingulate cortex. Active neurofeedback training and enhanced connectivity in cortical midline regions also tracked with improved reality monitoring accuracy following scanning for recollection of the source of imagined information, a behavioural effect not observed in the Sham group and not generalising to item recognition memory. This suggests a link between activity and connectivity changes within the paracingulate region and an improvement in reality monitoring for the source of internally generated information. Furthermore, the improvement in reality monitoring accuracy following Active neurofeedback was specific for recognition of the source of internally-generated (Imagined) stimuli. No significant improvement was observed in either group for stimuli which had previously been presented as Seen, nor for discrimination of the source of information which had earlier been spoken by the Subject or Researcher. Notably, there was also no improvement in accuracy for item recognition memory, suggesting that the effect of the Active neurofeedback training was not to enhance memory retrieval overall but was specific to improved recollection of the source of Imagined information. This finding is thus consistent with previous work associating activity within the PFC with recollection of the context in which previous events have occurred (33–35) and with medial PFC in particular associated with recognition of the source of internally generated information (6, 35, 36).

Reassuringly, the change in peak functional activity following Active neurofeedback training was observed within the mean paracingulate mask based on all 39 participants. It was measured in an mPFC ROI based on a brain region previously associated both with dysfunctional activity in schizophrenia and reality monitoring accuracy in healthy individuals (25, 32). The anterior location of this ROI within the frontal pole is consistent with imaging research findings on reality monitoring more generally (13). However, in Study 1, we found that the peak of the mPFC/paracingulate reality monitoring interaction for functional connectivity between patients with schizophrenia and healthy control groups (Study 1) lay more posterior to this suggesting a possible separation between the peak of reality monitoring activity and the peak of associated functional connectivity within the mPFC. This potential functional dissociation is clearly interesting and a focus for follow-up investigation. Inspection of the peak functional activity pattern within paracingulate cortex showed sequentially decreasing deactivation relative to the baseline in the Active group across the three feedback runs (Figure 3B). As the contrast used in the reality monitoring functional analysis was feedback > rest, deactivation within the paracingulate mask is consistent with more general DMN deactivation during task related activity (37). A sequential reduction in the scale of deactivation over the three neurofeedback runs might thus indicate increased self-referential processing as participants in the Active group became more successful at increasing their brain activity within the paracingulate mask. A similar pattern of reduced deactivation within the mPFC and DMN more broadly during reality monitoring was also described previously (4).

### Limitations

In Study 1, we restricted our participants to healthy controls and patients with schizophrenia with hallucinations and due to difficulties identifying suitable participants for recruitment, we did not include a third group of patients with no life-time experience of hallucinations. As such, we cannot be sure that our finding linking paracingulate functional connectivity with reality monitoring impairment also extends to the association between paracingulate morphology and hallucinations in schizophrenia (10, 11). While much previous work has identified a reality monitoring impairment associated with hallucinations in schizophrenia (13), this point provides a focus for replication and extension of the current study. We also included no test of recognition memory in Study 1, and thus cannot confirm that the reality monitoring impairment we observed was specific to recollection of the source of Imagined information. Indeed, reality monitoring impairment was also observed for recollection of the source of words spoken by the researcher suggesting that there may have been a general memory deficit across the patients in our sample. However, previous work has shown that impairments in reality monitoring in patients with schizophrenia exist even in the absence of recognition memory deficits, and also that the enhanced externalisation bias which underlies impaired reality monitoring for Imagined items in schizophrenia is associated with the experience of hallucinations (13). Thus, the impairment in reality monitoring accuracy for Imagined items observed in this sample of patients may well have been a factor contributing to their hallucinatory experience (3, 13).

A further limitation of our work were the small sample sizes suggesting that the results should be taken as preliminary. In Study 2, we report an effect of fMRI neurofeedback training on reality monitoring for Imagined information in the Active group, but not the Sham group, but the interaction failed to reach significance (p = .109). We also reported differential functional connectivity following fMRI neurofeedback training between the paracingulate seed region and the precuneus in the Active group, but again, although suggesting a strong trend effect, the interaction with the Sham group failed to reach significance (p = .07). Finally, we found no significant correlation between the change in reality monitoring score for Imagined items, and the change in functional connectivity following Active neurofeedback training which would support our arguments that neurofeedback training acts to enhance reality monitoring ability by increasing connectivity within the highlighted medial cortical network. In each case, these finding may be explained by low power, and should be investigated in a replication study with a larger sample. Greater effect sizes and possibly longer term effects might also be obtained from neurofeedback using more than one training session (31).

### Conclusions

Our results suggest the presence of a processing network involving paracingulate cortex subserving reality monitoring in healthy individuals, while patients with schizophrenia appear to utilise a distinct and more lateral network. This may explain sub-optimal reality monitoring performance in patients, and may contribute to their experience of hallucinations. Activity and connectivity in the reality monitoring network in healthy individuals can be modulated by Active fMRI neurofeedback training, which tracks with in an improvement in reality monitoring accuracy for self-generated information. The involvement of midline structures (paracingulate and precuneus cortex) in reality monitoring is consistent with earlier work focusing on the DMN in self-referential processing generally (9), and on reality monitoring in particular (4). Together, the above findings suggest that neurofeedback training might have value as a treatment for hallucinations in schizophrenia by improving reality monitoring ability through alteration of the underlying functional networks. Recent fMRI neurofeedback proof of concept studies in schizophrenia add support to this proposal (30, 38). We suggest further work to focus on replication of these Studies in a larger sample and their extension to include a schizophrenia patient population comprising individuals both with and without the experience of hallucinations.

## Methods

### Study 1

#### Participants

Thirty-nine right-handed participants were recruited, comprising 19 patients (mean age = 36.9 years, SD = 7.1 years, 1 female), who met the DSM-5 criteria for schizophrenia (verified by clinical assessment through the Mini International Neuropsychiatric Interview (39) and 20 healthy controls (mean age = 33.4 years, SD = 8.0 years, 2 females) as previously detailed (12). One patient participant who had been recruited and whose data was included in a previous study was excluded from this analysis as they had no lifetime experience of hallucinations. Written informed consent was obtained from participants in a manner approved by the UK National Research Ethics Service. The patient and control groups were matched on age [t(37) = 1.641, n.s.], sex (χ^2^ = .003, n.s.) and verbal IQ [as measured by the National Adult Reading Test (40); t(37) = 1.510, n.s.]. The groups differed on fluid intelligence [as measured by the Raven’s Advanced Progressive Matrices (41); t(37) = 2.850, p = .007], and Launay Slade measures of hallucinations [see Exp. 2 for test details, t(37) = 4.330, p < .000]. All patients were receiving antipsychotic medication but none were on drug regimens that included typical antipsychotics, anticholinergics or benzodiazepines, with a mean time on medication of 12.63 years (SD = 4.71 years).

#### Study Protocol

Participants performed a reality monitoring task and a resting state fMRI scan in a single scanning session. Functional and behavioural analysis of the reality monitoring data, alongside additional results from this study and full methodological details can be found in (12).

#### Reality Monitoring Task

During each study trial the word-pair was shown, either complete (Seen trials) or with only the first letter of the second word provided such that the second word needed to be self-generated (Imagined trials; See Figure 5). The subject or researcher then had 2.5 seconds to read aloud the word-pair, completing it as necessary for Imagined trials. Each study phase was followed by its corresponding test phase, during which subjects were presented with the first word from each word pair in the study phase and were required to indicate if ‘the accompanying word was Seen or Imagined?’ Participants had 4.5 seconds in which to respond. The order of presentation of sub-blocks in the test phase alternated across the six full blocks of the task and was counterbalanced across participants. A separate source monitoring condition was also tested in which the subjects were tasked with recalling whether they or the researcher had read allowed the word-pair (this data is not included here but has been reported elsewhere: (12)). The task comprised six separate study and test blocks, with 24 word-pair stimuli in each study phase, with word-pairs assigned to each of the conditions counterbalanced across participants.

**Figure 5.**
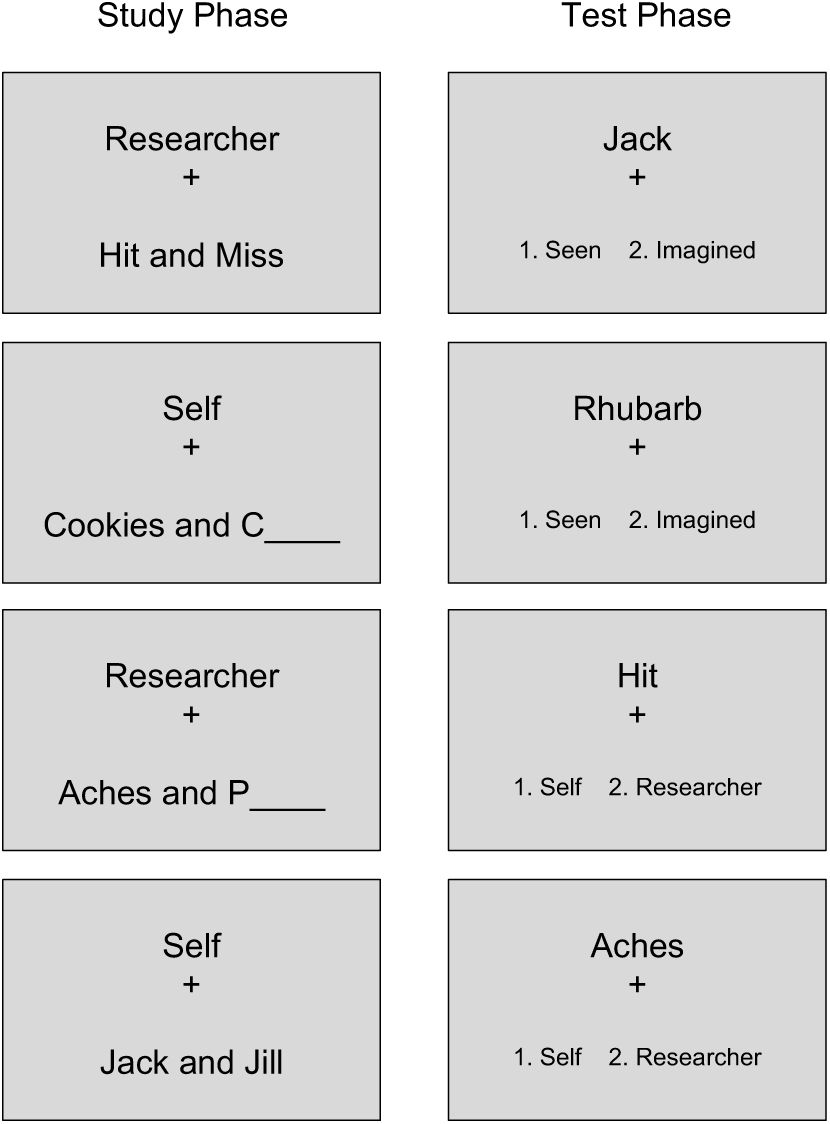
Stimuli used in the study and test phases of the Reality Monitoring Tasks in Study 1. Note: In a 2 × 2 design, either the subject or researcher spoke aloud the word-pair stimuli, which were presented either complete (Seen) or incomplete (requiring the second word to be Imagined). Subjects were then presented at test with the first word of a word-pair, and asked to respond as to whether the accompanying word had been Seen or Imagined, or whether they or the researcher had read aloud the word-pair

Post testing, the reality monitoring accuracy data was analysed using a group x task condition ANOVA and between subjects t-tests.

#### MRI acquisition

Imaging data were acquired using a 3T Siemens TIM Trio scanner at the Wolfson Brain Imaging Centre on the Addenbrookes Biomedical Campus, Cambridge, using a 32-channel head coil. Each session began with a localizer scan, followed by a structural T1-weighted MP RAGE image with spatial resolution of 1 mm^3^ in plane resolution, 256 × 240 × 176 slices. Resting state images were acquired following collection of behavioural and functional task data, using a full-brain, T2* weighted, BOLD-sensitive gradient echo planar sequence: TR = 2.14 seconds, TE = 30 mseconds, flip angle 78°, field of view 192mm × 192mm, 36 slices, 2mm slice thickness and in plane resolution of 3mm × 3mm. 280 volumes were collected during the resting state scan.

#### Resting state functional connectivity analyses

Resting state data were analysed with FMRIB Software Library (FSL) (42). Volume reorientation and head motion correction was performed using MCFLIRT software (43) with rigid body transformation. Brain extraction was then undertaken on both the T1-weighted images and EPI motion corrected scans using BET (44) with the f parameter set to 0.5, followed by visual inspection of the images to ensure appropriate extraction of the brain. The images were smoothed using an isotropic 6mm full-width half maximum Gaussian kernel (twice the voxel size of the images) (45), with a high-pass linear filter cut off of 100 mHz. Co-registration and normalization to standard MNI template were undertaken using FLIRT (43) by co-registration of the mean functional image to the T1-weighted brain extracted image, then applying the transformation between the T1 anatomical images to MNI space (trilinear interpolation) to the pre-filtered functional sequence. The final resolution of the functional image, after normalization resampling, was set to 4mm.

The pre-processed resting-state fMRI data was analysed using Multivariate Exploratory Linear Optimized Decomposition into Independent Components 3.0 (MELODIC)(42). The multiple 4D data sets were decomposed into their distinct spatial and temporal components using ICA, and temporarily concatenated (42). A single 2D analytical run was undertaken on the concatenated data matrix obtained by stacking the 2D data matrices of every dataset for all subjects in the group, with the number of Independent Components set manually to 30 (46–49).

In order to separate noise components from the underlying resting-state networks, Independent Components were tested for their correlation to labelled networks from the 7-network cortical parcellation of Yeo et al. (18). Given our *a priori* hypotheses, we focused our exploratory analysis of the functional connectivity data on the DMN and SSN (18). The two Independent Components used in the functional connectivity analysis were chosen after establishing a threshold of r-value > .2 of correlation as recommended in FSL (42) with the DMN and SSN reference networks (18). The actual correlations achieved were well in excess of this threshold. The Independent components used and their comparison to the reference networks are shown in Supplementary Materials Figure S1.

The selected Independent Components were submitted to second level analysis. Functional connectivity group effects between schizophrenia and controls groups were tested using independent t-test and dual regression. Thereafter, reality monitoring accuracy scores for Imagined items were added as a variable of interest in the general linear models to investigate group interactions between functional connectivity and reality monitoring accuracy across networks. Statistical group differences were tested using non-parametric permutation testing, with threshold-free cluster enhancement (50). Functional connectivity results are reported for p < .05 familywise-error (FWE) threshold corrected for multiple comparisons across voxels, and p < .025 with Bonferroni correction for multiple comparisons across the two Independent Components (networks).

### Study 2

#### Participants

39 healthy individuals (males = 15) participated in a single-blind randomised controlled Study. None of these individuals took part in Study 1. Participants were recruited from the University of Roehampton, Royal Holloway University of London and from the general public using adverts on social media. The University of Roehampton Ethics Committee gave ethical approval and all participants gave written informed consent prior to taking part in the study. The mean age of participants was 21.9 years (SD = 3.0 years) and there were 36 right-handed and 3 left-handed participants. Participants had no prior neurological or medical illness and were not using any psychiatric medication. 21 participants were randomly assigned to the ‘Active’ neurofeedback condition targeting the mPFC, and 18 participants to the ‘Sham’ neurofeedback condition, with no significant differences between the groups in terms of age [t(37) = 1.154, n.s.], sex (χ^2^ = .003, n.s.), or handedness (χ^2^ = 2.786, n.s.).

#### Study Protocol

Participants underwent questionnaire assessment, offline reality monitoring testing and MRI scanning in a single three-hour session. Details of the study protocol are given in Figure 6.

**Figure 6.**
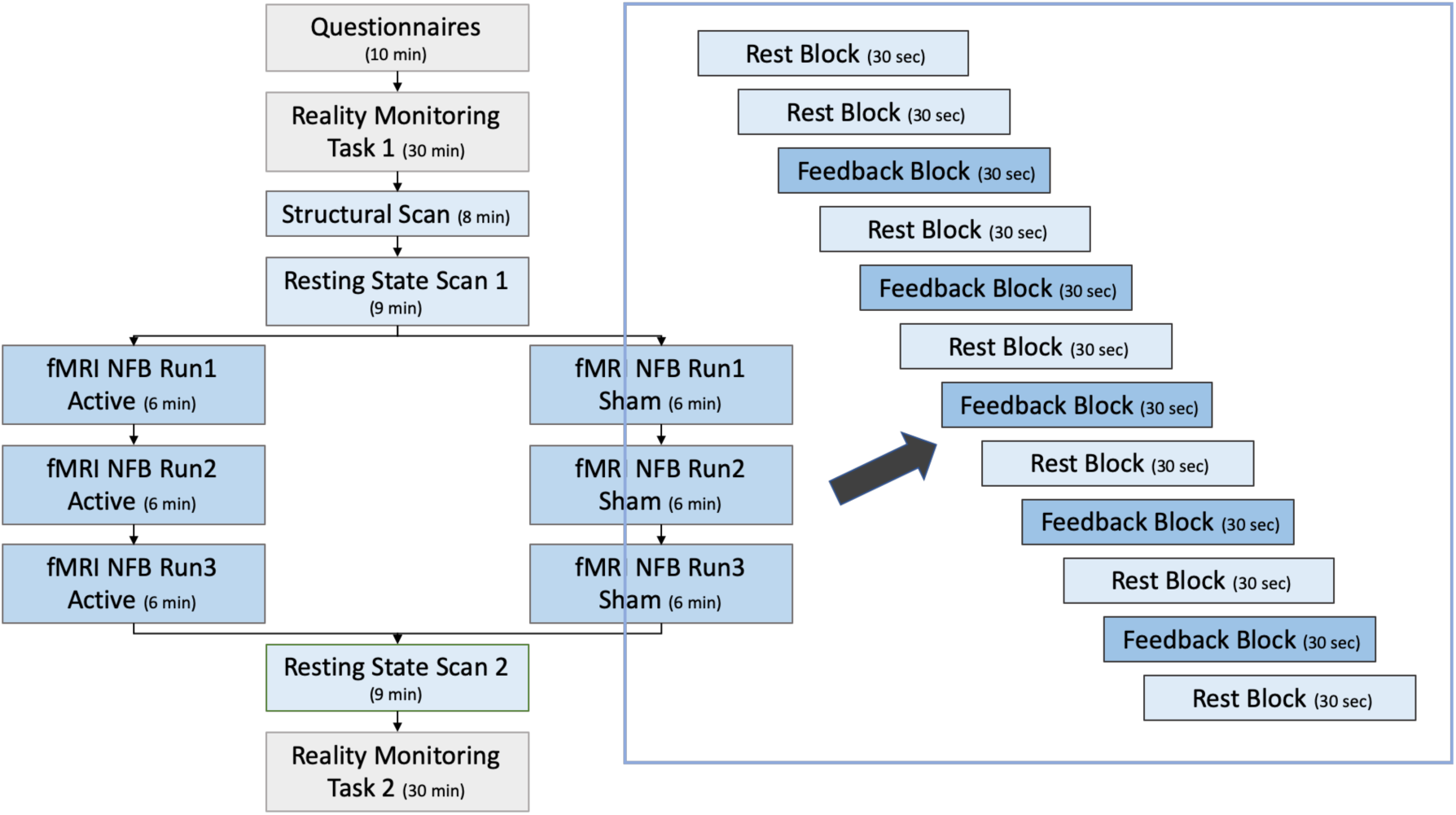
Study 2 Study Protocol. Participants were randomly allocated to Active and Sham groups, but aside from neurofeedback, experienced identical protocols. The right-hand side of the image indicates the neurofeedback run in detail. Blue text-boxes indicate scanned sessions, Grey boxes indicate off-line measures. Note: NFB = neurofeedback. An extra rest session is included at the start of each imaging run in order to establish a longer baseline for the neurofeedback contrast.

#### Assessment for schizotypy and proneness to hallucinations

Individuals’ proneness to hallucinations and schizotypy was assessed by self-report using a written questionnaire prior to scanning, to ensure that the groups were matched on trait measures previously related to reality monitoring and hallucinations. Hallucinations proneness was assessed using the Revised Launay-Slade Hallucination Scale (LSHS-R) (51, 52). Schizotypy was assessed using the Brief O-LIFE scale (OLIFE-B), a 30-item shortened version of the original 104-item Oxford-Liverpool Inventory of Feelings and Experiences (53, 54). Active and Sham neurofeedback groups did not differ in terms of O-LIFE measures of schizotypy [t(37) = .695, n.s.] or Launay Slade measures of hallucinations [t(37) = 1.206, n.s.).

#### Reality Monitoring Task

A word-pair reality monitoring task was used, similar in design to that described for Study 1. Two versions of the task were used, one undertaken prior to the participant entering the scanner, and the second immediately after the participant exited the scanner, with new word-pairs in the second task. Unlike in Study 1, these versions of the task included an additional recognition memory Old/New condition, whereby additional new words not included in the study phase were added in to the test phase and the participants given the additional option of responding ‘New’ if they did not previously remember seeing the word. Each task comprised five separate study and test blocks, with 24 word-pair stimuli in each study phase, 12 additional new words included in the test phase and all stimuli counterbalanced across participants.

Old/New recognition memory accuracy was calculated as the adjusted item recognition score (hits minus false alarms, with hits being defined as the proportion of words correctly recognised as previously seen and false alarms the proportion of New words incorrectly endorsed as Old). Reality monitoring accuracy was calculated as the number of accurate source responses divided by the number of correct responses recognising an item as Old.

#### MRI acquisition

All MRI scans were acquired on a 3 Tesla Siemens Magnetom TIM Trio scanner using a 32-chanel head coil at the Combined Universities Brain Imaging Centre at Royal Holloway, University of London (CUBIC; http://www.cubic.rhul.ac.uk). Each participant underwent an anatomical scan which comprised a T1-weighted Magnetization Prepared Rapid Acquisition Gradient Echo (MP RAGE) image (1mm^3^ resolution, in plane resolution 256 x 256 x 176 slices, acquisition time approximately 5 minutes). Three separate fMRI neurofeedback runs were acquired in each participant comprising of five feedback vs. seven rest blocks each lasting 30 seconds (Figure 6). Resting state fMRI scans were acquired in all participants before and after the acquisition of the 3 fMRI neurofeedback training runs. All functional resting state and neurofeedback scans were acquired using echo-planar image sequences: TR = 2 seconds, TE = 40 mseconds, 28 slices, 4mm slice thickness, in-plane resolution 3mm x 3mm. As with the resting state scans in Study 1, subjects were instructed to rest with their eyes closed, and to try not to fall asleep.

#### Anatomical Localiser

The paracingulate target region for fMRI neurofeedback was delineated anatomically in all participants using their T1 weighted anatomical scan. An investigator trained to recognise the morphology and anatomy of the region (JG) manually delineated an anatomical region along the bilateral paracingulate sulcus, using tools in BrainVoyager (Brain Innovation, Maastricht, Netherlands). The extent of the combined binary mask across all participants is shown in Figure 2. The PCS target regions were then transferred into a Turbo-BrainVoyager (version 4) file format and used to define the neurofeedback target region (VOI) on echo-planar images during the neurofeedback training runs.

#### fMRI Neurofeedback

FMRI Neurofeedback was administered over three x 6 minute runs during a single visit using Turbo-BrainVoyager with each run composed of Feedback and Rest blocks (see Figure 6). Reconstructed DICOM images were directly transferred from the MRI scanner via a secure data transfer protocol to an analysis computer where Turbo-BrainVoyager was installed. Pre-processing was performed in real time on all transferred images, including Gaussian spatial smoothing with a kernel of 4mm full width half maximum, and motion correction. The functional data was registered to the anatomical scan acquired at the beginning of the scanning session.

Participants received either ‘Active’ veridical (based on the fMRI neurofeedback signal from the paracingulate VOI) or Sham signal during Feedback blocks via a visual thermometer gauge interface (Figure 3D). Participants were instructed to move the thermometer up during Feedback blocks, so that all cells in the gauge were turned grey (achieved by up-regulation of the BOLD signal in the VOI). Participants were instructed to relax during Rest blocks. No specific direction or instructions were given to participants regarding how to self-regulate their neurofeedback signal (55) but participants were told to allow 5 to 7 seconds for their efforts to result in a change in the gauge (to allow for the haemodynamic response). During Feedback blocks, a continuous signal from the VOI target area was displayed via the visual gauge and updated for every scan volume (TR = 2 seconds). The feedback signal was calculated using a real-time general linear model based on a single predictor for the Feedback/Rest onsets function convolved with the haemodynamic reference function. The Sham feedback signal was based on a saved pattern of randomised activity at a similar level of intensity to active feedback (provided by Turbo-BrainVoyager technical support), but derived from no specific brain region. Changes in amplitude of the thermometer were indicated in terms of the percentage signal change, calculated as the current signal value compared with the average value determined from the immediately preceding rest block (56). The thermometer gauge was scaled with a maximum value of 0.5%, and gradations of 0.05%, chosen to match previous successful neurofeedback studies (29, 57). The thermometer remained visible during rest blocks, but no feedback was given. Thereafter, a change in colour of the top box on the thermometer gauge from red to white indicating a switch from a Rest to a Feedback block, simultaneously accompanied by two second presentation of the words ‘Rest’ (and vice versa with the words ‘Move Thermometer’).

#### fMRI data analyses

Functional data were analysed using SPM12 (http://www.fil.ion.ucl.ac.uk/spm). Functional volumes were spatially realigned to the first image of the first series and volumes normalised against the MNI reference brain using tri-linear interpolation, and smoothed with an isotropic 8mm full-width half-maximum Gaussian kernel. First level modelling was undertaken using a block-related design with separate regressors coding for the onsets and durations of Feedback and Rest blocks. These together with the six regressors coding movement parameters, comprised the full model for each neurofeedback run. The data and model were high-pass filtered to a cut off of 1/ 128 Hz.

A simple contrasts of interest analysis was performed on individual subject data at the first level comparing Feedback > Rest blocks. Second-level modelling was then performed on the combined single-subject contrasts to produce random-effects group analyses separately for each run in the Active and Sham groups, and to enable random-effects between-group analyses to test the interaction between group (Active vs. Sham) and neurofeedback training run (Run1, Run2, Run3). We then tested our a priori hypothesis that subjects receiving active neurofeedback would be able to up-regulate activity in the VOI across 3 Active neurofeedback runs during the single scanner visit. To do this, we examined this interaction term within a paracingulate ROI defined as an 8mm radius sphere chosen to match the width of the smoothing kernel used in the fMRI preprocessing (and consistent with our previous functional activity and morphological research methods on reality monitoring and PCS morphology (10, 12)). To avoid circular inference, the sphere was centred on *a priori* coordinates of a PFC brain region that is associated with reality monitoring in healthy subjects (25) and shows dysfunctional activity in schizophrenia [Talairach: 14, 47, −1, Blue sphere in Figure 2]; (32). This sphere overlapped with the combined binary VOI mask across all subjects (Figure 2). A small volume correction was used on the whole-brain Feedback > Rest first level contrasts with a family-wise error corrected voxel-wise height threshold of p < 0.05.

#### Resting state functional connectivity analyses

Resting state fMRI data (pre and post neurofeedback) were pre-processed as described in Study 1. The aim of our analysis was to compare differences post > pre neurofeedback training between groups in reality monitoring-related functional connectivity from paracingulate cortex. Therefore, for each subject and each session, we created a mask of the seed region identified as the paracingulate reality monitoring x functional connectivity interaction peak from Study 1 [−14, 38, 20]. Being so frontal within the brain, paracingulate cortex is especially susceptible to field distortion. Therefore, to improve reproducibility in the connectivity analysis across subjects (see(58, 59)) we used a relatively large sphere of 10 mm radius as seed region (Green sphere in Figure 2). For each subject in each session, we then extracted the average time series defined by our binary mask and calculated the seed-based correlation by means of dual regression.

Pre-training spatial maps for each subject were first contrasted with post-training maps (i.e. post > pre neurofeedback), and then input into a higher-level analysis to investigate group differences (Active vs. Sham) in functional connectivity related to the neurofeedback training. Statistical group differences were tested using non-parametric permutation testing, with threshold-free cluster enhancement (50). Finally, we examined the interaction between changes in reality monitoring accuracy for Imagined items and changes in functional connectivity (pre vs. post neurofeedback) within the Active neurofeedback group. Functional connectivity results are reported for p < .05 FWE threshold corrected for multiple comparisons across voxels.

## Acknowledgments

JRG and CF were supported by a Wellcome Trust Collaborative Award (WT108720).

## Supplementary Materials

**Figure S1.**
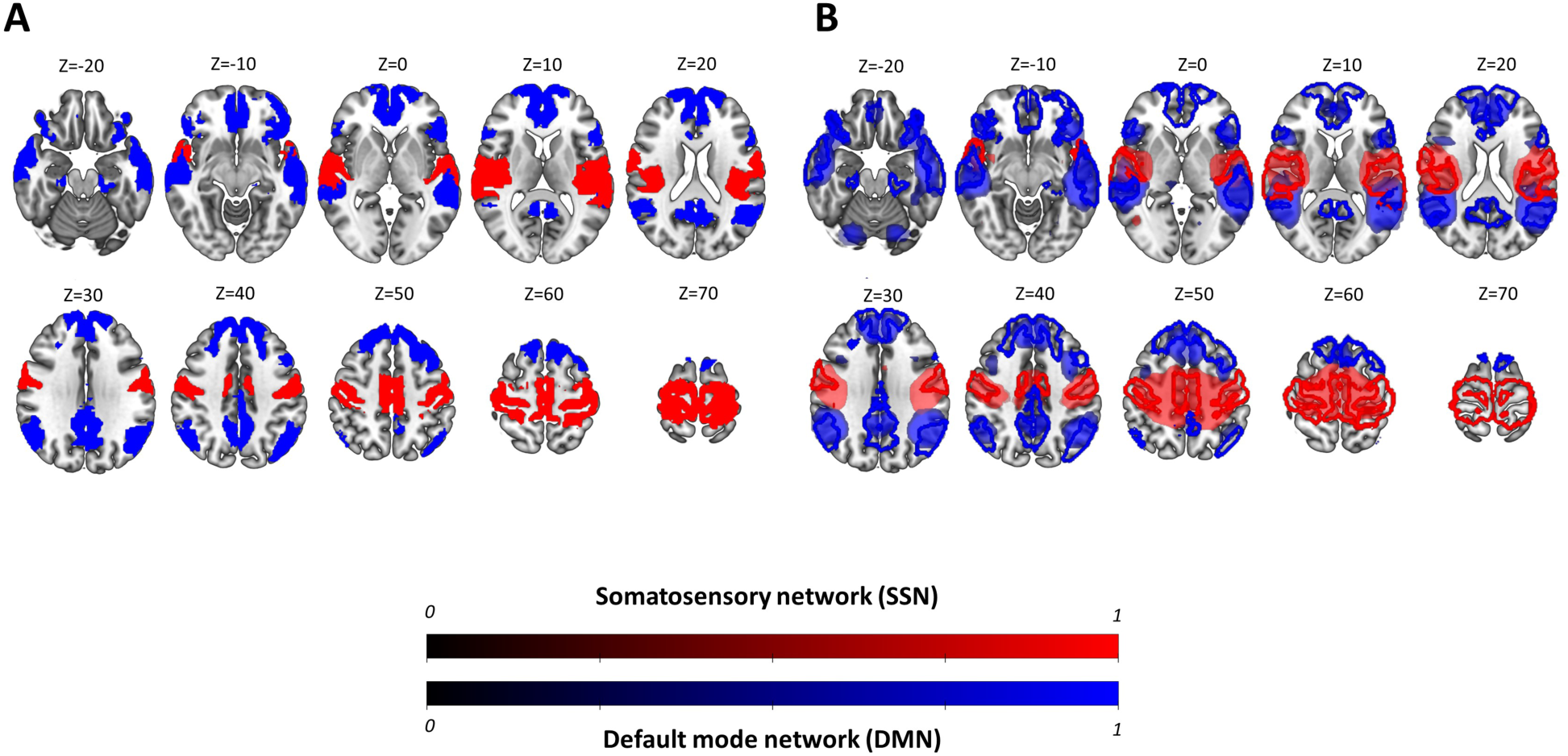
Study 1 (A) Somatosensory (Red; SSN) and Default Mode (Blue, DMN) networks from the 7-networks parcellation of the human cortex by Yeo et al. (2011); (B) The Independent Components selected for analysis from the resting state data in Study 1 are shown to highlight the overlap with these networks. Pearson’s correlation coefficients were as follows: SSN r = 0.50; DMN r = 0.49.

**Figure S2.**
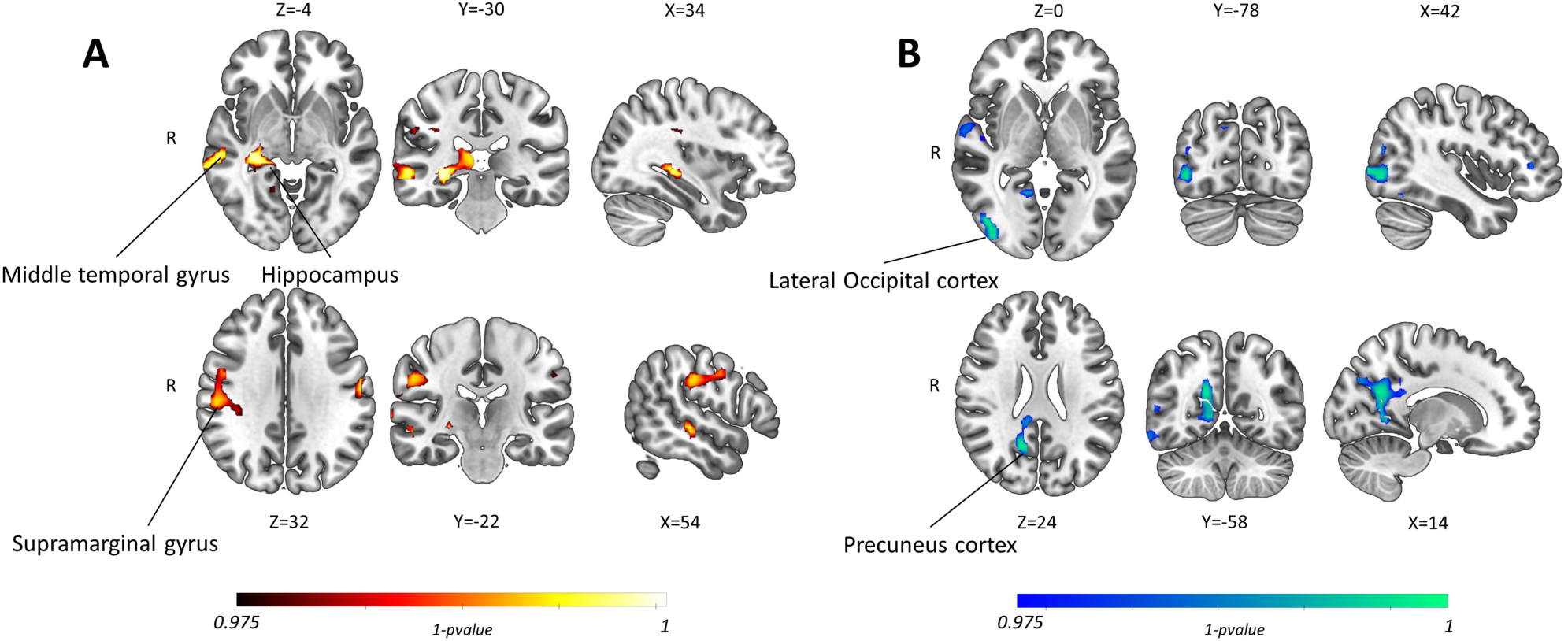
Study 1. Somatosensory Network (SSN) functional connectivity changes in Schizophrenia relative to healthy controls showing (A) Increase (Red/Yellow), and (B) Decrease (Blue/Green) in SSN functional connectivity in Schizophrenia patients relative to healthy controls.

**Figure S3.**
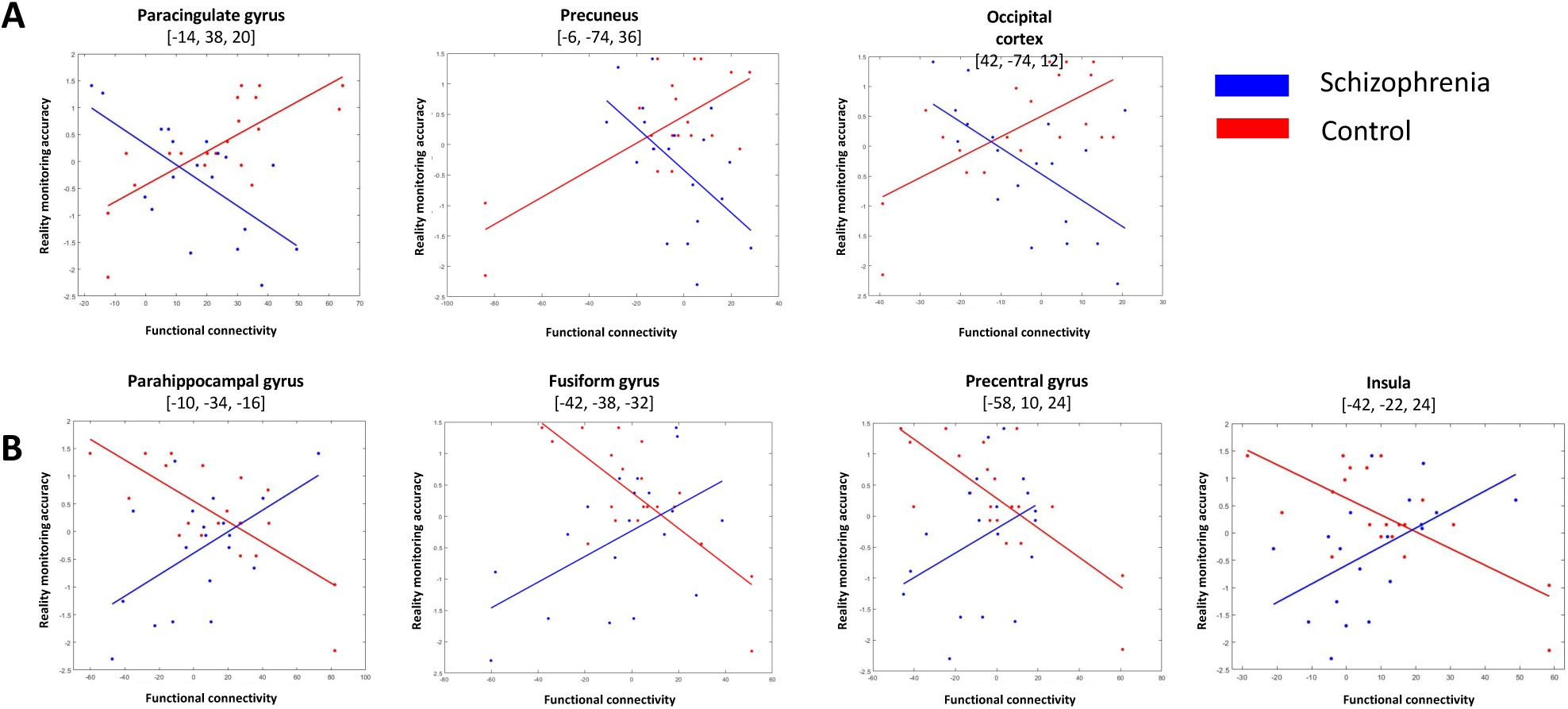
Study 1. Scatter plots showing positive (A, upper row) and negative (B, lower row) association between reality monitoring accuracy for Imagined items and functional connectivity parameter estimates from observed peaks for Schizophrenia (Blue) and Healthy Control (Red) Participants

**Table S1.** Study 1. Differences in functional connectivity between Patients with Schizophrenia and Healthy Controls for the Default Mode and Somatosensory networks. p-value of the peak coordinates and anatomical labelling are shown. Where the Cluster comment is ‘none’, no clusters survived the Bonferroni corrected threshold of p < .025. Nb. Unless otherwise stated, all coordinates are in MNI space.

| IC Network & contrast | Cluster | p value | Peak coordinates |  |  | Extent of Coverage |
| --- | --- | --- | --- | --- | --- | --- |
|  |  |  | x | y | z |  |
| **Default Mode Network** |  |  |  |  |  |  |
| Schizophrenia > Controls | None |  |  |  |  |  |
| Controls > Schizophrenia | None |  |  |  |  |  |
| **Somatosensory Network** |  |  |  |  |  |  |
| Schizophrenia > Controls | 1 | .001 | 34 | -30 | -4 | Hippocampus, Lingual Gyrus, Parahippocampal Gyrus, Thalamus, Intracalcarine Cortex, Precuneus |
|  |  | .003 | 30 | -42 | 0 |  |
|  |  | .008 | 14 | -30 | 12 |  |
|  |  | .012 | 10 | -58 | 0 |  |
|  | 2 | .003 | 58 | -30 | -4 | Middle Temporal Gyrus, Superior Temporal Gyrus, Supramarginal Gyrus, Parietal Operculum Cortex, Planum Temporale, Postcentral Gyrus, Central Opercular Cortex, Angular Gyrus |
|  |  | .005 | 66 | -34 | -4 |  |
|  |  | .012 | 70 | -18 | 4 |  |
|  |  | .012 | 70 | -14 | 12 |  |
|  |  | .018 | 70 | -46 | 16 |  |
|  | 3 | .009 | 54 | -22 | 32 | Supramarginal Gyrus, Postcentral Gyrus, Parietal Operculum Cortex, Precentral Gyrus, Middle Frontal Gyrus |
|  |  | .012 | 50 | -10 | 36 |  |
|  |  | .013 | 58 | -2 | 0 |  |
|  |  | .019 | 38 | -30 | 32 |  |
|  | 4 | .002 | -62 | -14 | 24 | Postcentral Gyrus, Supramarginal Gyrus |
|  | 5 | .002 | -18 | 58 | 4 | Frontal Pole |
|  |  | .003 | -2 | 62 | 8 | Frontal Pole, Superior Frontal Gyrus |
|  | 6 | .020 | -6 | -30 | 0 | Thalamus |
| Controls > Schizophrenia | 1 | .004 | 42 | -78 | 0 | Lateral Occipital Cortex, Occipital Pole, Inferior Temporal Gyrus, Fusiform Gyrus, Middle Temporal Gyrus, Angular Gyrus |
|  |  | .006 | 42 | -90 | 0 |  |
|  |  | .009 | 50 | -74 | 16 |  |
|  |  | .017 | 50 | -66 | -12 |  |
|  |  | .020 | 58 | -58 | 12 |  |
|  | 2 | .004 | 14 | -58 | 24 | Precuneus, Supracalcarine Cortex, Lingual Gyrus, Intracalcarine Cortex |
|  |  | .009 | 10 | -58 | 4 |  |
|  |  | .009 | 14 | -58 | 12 |  |
|  | 3 | .010 | 26 | -6 | 4 | Right Putamen, Right Pallidum |
|  | 4 | .021 | 10 | -38 | 24 | Cingulate Gyrus |
|  | 5 | .023 | 14 | -78 | 36 | Cuneal Cortex, Precuneus, Lateral Occipital Cortex |
|  | 6 | .024 | 66 | -54 | -16 | Inferior Temporal Gyrus, Middle Temporal Gyrus |
|  | 7 | .022 | 42 | 38 | 4 | Frontal Pole, Inferior Frontal Gyrus |
|  | 8 | .017 | 58 | -6 | 2 | Postcentral Gyrus, Precentral Gyrus |

**Table S2.** Study 1. Interaction effects between group (Schizophrenia vs Control) and reality monitoring (‘RM’) accuracy for Imagined items for functional connectivity in the Default Mode and Somatosensory Networks. Where the comment is ‘None’, no clusters survived the Bonferroni corrected threshold of p < .025. Scatter plots for these interaction effects are shown in Figure S3.

| Network & Interaction | Cluster | p value | Peak coordinates |  |  | Extent of Coverage |
| --- | --- | --- | --- | --- | --- | --- |
|  |  |  | x | y | z |  |
| **Default Mode Network** |  |  |  |  |  |  |
| Schizophrenia vs. Control x RM Accuracy | 1 | .001 | -14 | 38 | 20 | Paracingulate Gyrus, Cingulate Gyrus |
|  | 2 | .017 | 42 | -74 | 12 | Lateral occipital Cortex |
|  | 3 | .022 | -6 | -74 | 36 | Precuneus, Cuneal Cortex |
| Control vs. Schizophrenia x RM Accuracy | 1 | .003 | 42 | -38 | -32 | Temporal Occipital Fusiform Cortex, Culmen |
|  | 2 | .004 | -10 | -34 | -16 | Parahippocampal Gyrus, Culmen |
|  | 3 | .013 | -42 | -22 | 24 | Central Opercular Cortex, Insula |
|  | 4 | .016 | -58 | 10 | 24 | Precentral Gyrus, Inferior Frontal Gyrus |
| **Somatosensory Network** | None |  |  |  |  |  |

**Table S3.** Study 2. Functional connectivity changes with paracingulate cortex, in the Active neurofeedback group following fMRI neurofeedback

| Cluster | p value | Peak coordinates |  |  | Extent of Coverage |
| --- | --- | --- | --- | --- | --- |
|  |  | x | y | z |  |
| 1 | .009 | -22 | -74 | 48 | Lateral Occipital Cortex, Precuneus, Superior Parietal Lobule, Supramarginal Gyrus |
|  | .009 | -34 | -58 | 48 |  |
|  | .010 | -50 | -34 | 48 |  |
|  | .012 | 58 | -34 | 48 |  |
|  | .013 | 34 | -66 | 48 |  |
|  | .013 | 18 | -74 | 64 |  |

